# Integrating T-cell receptor and transcriptome for large-scale single-cell immune profiling analysis

**DOI:** 10.1101/2021.06.24.449733

**Authors:** Felix Drost, Yang An, Lisa M Dratva, Rik GH Lindeboom, Muzlifah Haniffa, Sarah A Teichmann, Fabian Theis, Mohammad Lotfollahi, Benjamin Schubert

**Affiliations:** Computational Health Center, Helmholtz Munich, Neuherberg, Germany; School of Life Sciences Weihenstephan, Technical University of Munich, Munich, Germany; Wellcome Sanger Institute, Cambridge, UK; Department of Mathematics, Technical University of Munich, Munich, Germany; Department of Physics, Cavendish Laboratory, University of Cambridge, Cambridge, UK; Biosciences Institute, Newcastle University, Newcastle upon Tyne, UK

## Abstract

Recent advancements in single-cell immune profiling that enable the measurement of the transcriptome and T-cell receptor (TCR) sequences simultaneously have emerged as a promising approach to study immune responses at cellular resolution. Yet, combining these different types of information from multiple datasets into a joint representation is complicated by the unique characteristics of each modality and the technical effects between datasets. Here, we present *mvTCR*, a multimodal generative model to learn a unified representation across modalities and datasets for joint analysis of single-cell immune profiling data. We show that *mvTCR* allows the construction of large-scale and multimodal T-cell atlases by distilling modality-specific properties into a shared view, enabling unique and improved data analysis. Specifically, we demonstrated *mvTCR’s* potential by revealing and separating SARS-CoV-2-specific T-cell clusters from bystanders that would have been missed in individual unimodal data analysis. Finally, *mvTCR* can enable automated analysis of new datasets when combined with transfer-learning approaches.

Overall, *mvTCR* provides a principled solution for standard analysis tasks such as multimodal integration, clustering, specificity analysis, and batch correction for single-cell immune profiling data.

## Introduction

T cells are one of the critical components of the adaptive immune system. Their primary function is the detection of pathogens and control of the immune reactions through antigen recognition by their highly-diverse T-cell receptor (TCR). While recognizing antigens and immune signaling are well-researched individually, the interplay between T-cell function through the TCR and its phenotype remains largely unexplored. Recent findings have shown that T cells sharing the same TCR, so called clonotypes, express similar transcriptional phenotypes and distribute non-randomly across gene expression-based clusters [1, 2]. These findings indicate that antigen recognition through TCRs imprints specific transcriptional cell states shared across clonotypes making it necessary to jointly analyse TCR and transcriptomic information.

Paired measurements of TCR and transcriptome can be realized with modern single-cell multiomic sequencing techniques [3, 4], enabling the study of cell state and function, simultaneously [5–7]. However, as of now, both modalities are usually analyzed separately on the transcriptomic- and the TCR-level, potentially missing crucial interdependencies between the two modalities. Recent endeavors sought to integrate transcriptomics and TCR information. For example, Schattgen et al. used clonotype neighbor graph analysis (*CoNGA*) to detect correlations between TCR sequences and transcriptome [8]. T cell clones were identified that shared similar TCRs and gene expression profiles. Zhang et al. developed a Bayesian model called TCR functional landscape estimation supervised with scRNA-Seq analysis (*tessa*) to correlate both modalities and cluster T cell clones by their specificity [9]. While these methods incorporate both modalities for clustering, they do not provide an integrated representation for other downstream analyses, do not offer principled approaches to integrate multiple datasets, and scale only to small-size datasets hindering large-scale studies.

Furthermore, they resort to a clonotype-level approach, fusing cells with similar TCRs. However, cells from the same clonotype can have other characteristics on the transcriptomic level [10, 11]. This phenotypic differentiation of cells belonging to a clonotype during development or upon immune modulation is lost when reducing their cells to common gene expression profiles.

Here, we introduce *mvTCR* - a multi-view deep learning model for integrating TCR and transcriptome. *mvTCR* provides a cell-level embedding incorporating both modalities, seamlessly integrates into standard single-cell analysis workflow, and scales well to atlas-level analysis. We applied *mvTCR* to five T cell datasets to show that the shared representation offers a holistic view for immunological research. At the same time, we demonstrate that *mvTCR* preserves cell state and phenotype information better than other integration approaches. Furthermore, we demonstrated the method’s capability to reveal separate clusters of disease-specific and bystander T cells, which were unidentifiable in unimodal analysis on a recent SARS-CoV-2 single-cell dataset [7].

## Results

### Joint integration of TCR and gene expression through multiview variational autoencoders

*mvTCR* is a deep neural network, which follows the structure of a Variational Autoencoder (VAE) (Fig. 1a, **Methods** mvTCR). As input, the model receives the gene expression data 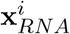 and the amino acid sequence of the TCR’s Complementary Determining Region 3 (CDR3) from the *α*-chain 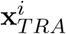 and *β*-chain 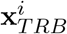 for each cell *i*. Following [13, 14], we employed a multi-layer perceptron to embed 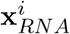 to a lower-dimensional representation 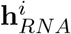. Due to its broad applicability on sequence data, we employed a transformer network to derive a representation of the TCR sequence 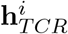. After encoding both modalities individually, a mixture module *M* was used to fuse both modalities into 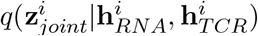 to derive a shared representation for each cell which can be used for various downstream analyses. We implemented three approaches to combine multiple modalities for *M*: Concatenation, Product-of-Experts (PoE) [15], and Mixture-of-Experts (MoE) [16]. The concatenation model simply combines both latent embedding 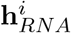 and 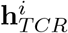 followed by an additional encoding network estimating the distribution of 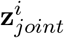. PoE and MoE first estimate sepa-rate marginal posterior distributions 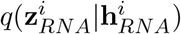 and 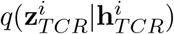, which were then fused via multiplication or addition to form the joint posterior distribution, respectively. As unimodal baselines, a TCR and a transcriptome model directly estimated the latent distribution from the 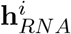 or 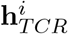 were additionally implemented. To train the VAE, similar decoding networks reconstructing 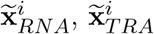, and 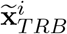 were used.

**Figure 1.**
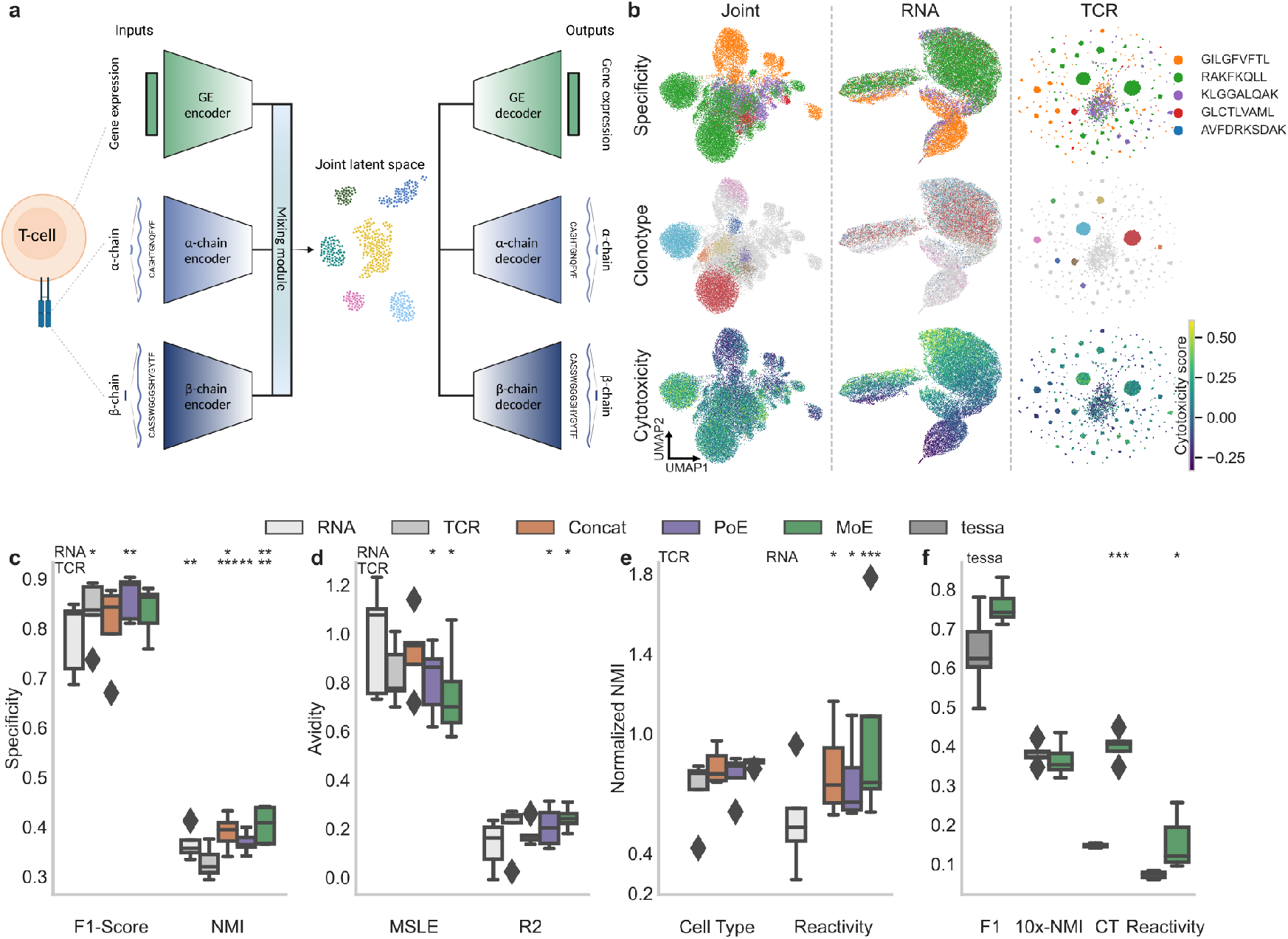
mvTCR jointly embeds TCR and transcriptome. **a**, Overview of the *mvTCR* architecture and the different mixture models. **b**, UMAP visualizations [12] of the 10x Genomics dataset (donor 2) comparing the embeddings of the multimodal (MoE) and unimodal (RNA, TCR) models colored by epitope specificity, ten largest clonotypes, and a cytotoxicity score as an example of cell state. **c-e** Comparison between the unimodal and all versions of the multimodal embeddings (Concatenation, PoE, MoE). Statistical significance to the corresponding unimodal embedding is calculated via one-sided, paired t-test (p-values: *<0.05, **<0.01, ***<0.001). The boxplot represents the quartiles and median line, while the whiskers extend to the full value range excluding outliers. **c**, Capturing of specificity by atlas-query prediction (weighted F1-Score) and clustering (Normalized Mutual Information) on the 10x dataset (donor 2). **d**, Avidity prediction measured by mean squared logarithmic error and R^2^ value on 10x dataset (donor 2). **e**, Conservation of modality-specific properties on the Fischer dataset by clustering (NMI, normalized by performance of underlying modality), i.e., cell type defined on transcriptomic level and reactivity towards a SARS-CoV-2 mix defined on a clonotype level. **f**, Comparison between MoE embedding trained only on gene expression and CDR3*β* and *tessa* [9] on the tasks defined in c and e.

### mvTCR incorporates knowledge from transcriptome and TCR

When analyzing T-cell repertoires, the antigen specificity of each cell is critical to provide the context for its cell state. The specificity of a T cell towards their cognate epitope is inherently determined by their individual TCR sequence with similar TCR recognizing similar epitopes. Additionally, cells with shared specificity express similar phenotypic characteristics. Therefore, we investigated to what extend a joint T cell embedding will benefit the preservation and prediction of antigen specificity. To this end, we used a dataset from 10x Genomics with binding annotations **s**^*i*^ of 44 epitopes for four donors (**Methods** Datasets). 68.2% of the T cells of donor 3 were specific to the CMV epitope *KLGGALQAK* and 26.0% did not express binding to any of the tested epitopes. 80.9% of donor 4 cells were considered non-binders. Hence, these donors were excluded during benchmarking due to the low amount of diverse annotation. For the remaining dataset, we separately evaluated *mvTCR* on T cells with specificity towards the eight most common epitopes for donor 1 with 9,639 cells (Supplementary Fig. 1) and donor 2 with 27,171 cells (Fig. 1b, Supplementary Fig. 2a). To observe the benefit of a multimodal embedding, the repertoire of both donors was embedded with uni- and multimodal versions of *mvTCR*. The unimodal embedding trained solely on **x**_*TCR*_ (Fig. 1b col.: TCR) was dominated by large clonotypes, which form separated clusters of distinct specificities. However, the embeddings of different clones did not follow a clear transcriptional pattern. The transcriptomic model trained on **x**_*RNA*_ (Fig. 1b col.: RNA) led to a more continuous representation, which formed several antigen-specific clusters. However, various subpopulations such as T cells binding to *GLCTLVAML* remained hidden. The multimodal models (Fig. 1b col.: MoE, Supplementary Fig. 2 row: Concatenation, PoE) revealed these clusters by conserving clonotype and cell state (e.g., Cytotoxicity Score, **Methods** Datasets), simultaneously (Fig. 1b).

### Antigen specificity is better captured in multimodal models

We then quantitatively evaluated *mvTCR’s* capability to capture antigen specificity on five random splits of the 10x dataset (Fig. 1c, Supplementary Fig. 3a, Supplementary Data 1, **Methods** Benchmarks). To this end, we first trained the models on a subset of all cells to build a reference atlas. Following, we simulated the annotation of novel T cells to their cognate epitope by mapping holdout cells into the reference atlas and predicting the cell’s antigen specificity via a k-Nearest-Neighbor (kNN) model. Note, that *mvTCR* was trained only on clonotypes which were not contained in the holdout set, to ensure unbiased performance estimation. All multimodal models outperformed the transcriptomic model (F1-Score, donor 1: 0.67, donor 2: 0.78) by 0.025 (Concatenation) to 0.07 (PoE) F1 points on donor 2 (Fig. 1c, first column) and 0.05 (Concatenation) to 0.15 (MoE) F1 points on donor 1 (Supplementary Fig. 3a, first column). Even though the TCR model visually showed a highly fractured embedding, it performed similar to the joint embeddings (F1-Score, donor 1: 0.735, donor 2: 0.835). Evidently, small sets of clones expressing the same specificity were grouped together locally in the TCR embedding, while global coherence was missing. From the multimodal models, PoE and MoE surpassed Concatenation for both donors.

Next, we evaluated the quality of the Leiden clusters [17] calculated on the different embedded spaces with regards to antigen specificity by Normalized Mutual Information (NMI) (Fig. 1c, Supplementary Fig. 3a, Supplementary Data 1). Here, most of the multimodal models statistically outperformed both TCR- and transcriptome-embedding across both donors (NMI, one-sided, paired t-test, p-values<0.05 for five out of six models compared to RNA models and four out of six compared to TCR models), while MoE surpassed all other models with a NMI of 0.58 for donor 1 and 0.40 for donor 2. Interestingly, the RNA based model showed significantly higher clustering scores than the TCR model for donor 2 (Fig. 1c, NMI, one-sided, paired t-test, p-value=0.007), even though it had significantly lower scores for kNN prediction (F1-Score, one-sided, paired t-test, p-value=0.029). This is in line with our previous observation, that the TCR embedding was highly fractured, while the RNA model formed a more continuous embedding space. Overall, the multimodal models captured the advantages of both modalities. They locally preserved antigen specificity, while globally grouping larger specificity clusters based on their similar transcriptomic profile.

In a next step, we investigated how informative the latent embedding was not only with regards to antigen specificity, but also to binding strength. To this end, we trained an additional neural network, that received the embedding as input to predict the counts of detected pMHC multimers as an approximate measure of avidity **a**^*i*^ (**Methods** Avidity Prediction, Supplementary Data 1). For donor 2, the multimodal embedding proved to be beneficial (Fig. 1d). Similar to the previous kNN classification, Concatenation (Mean Squared Logarithmic Error (MSLE): 0.930, Coefficient of determination (R^2^): 0.179) was the weakest of the three multimodal versions, while PoE (MSLE: 0.811, R^2^: 0.209) and MoE (MSLE: 0.756, R^2^: 0.244) outperformed both unimodal embeddings (**Methods** Benchmarks). However, the prediction failed for donor 1 on all models (Supplementary Fig. 3b, all R^2^<0). The prediction was heavily biased towards the epitopes *IVTDFSVIK* and *AVFDRKSDAK* depending on whether the training or test set contained large clonotypes, which showed 81.7% and 75.4% of all bindings for these epitopes, respectively. Summarizing, we overall demonstrated that multimodal embeddings were more informative of antigen specificity than models trained on either the TCR or the transcriptome alone.

### The joint representation preserves cellular heterogeneity

Besides functional aspects, multimodal embeddings further need to conserve modality-specific characteristics such as cell type and clonotype. To test this, we trained *mvTCR* on five different subsamples of the dataset described in Fischer et al. [6] (Supplementary Data 1). This dataset contained 6,713 T cells of two SARS-CoV-2 infected patients with annotated T cell subtypes defined on the transcriptome (**Methods** Datasets). Additional, clonotypes were identified via reverse phenotyping, which were able to recognize a mix of disease-specific peptides. Ideally, the shared embedding should conserve these modality-specific annotations to a large degree during clustering. Therefore, we reported the NMI of the models normalized to the score obtained by the respective modality, on which the annotation was defined (Fig. 1e). This provided us with an estimate of how well modality-specific characteristics were retained. All multimodal models clustered the cell types better than the TCR-model (72.1%) reaching up to 85.7% (MoE) of the transcriptome model’s score. For clustering based on reactivity, all multimodal models significantly outperformed the transcriptome version (NMI, paired, one-sided t-test, p-values<0.05 for Concatenation and PoE, p-value=0.0004 for MoE), while the MoE-version achieved on average almost identical performance (99.1%) as the TCR-model. Hence, we concluded, that both TCR- and gene expression-specific characteristics are represented to a high degree in the shared embedding of *mvTCR*. Based on its high performance in most settings *mvTCR*’s MoE module was used for all following analyses.

Finally, we compared *mvTCR* against the weighted embedding derived by *tessa* for predicting specificity and conserving modality-specific annotation (Fig. 1f, Supplementary Fig. 3c) (**Methods** Benchmarks). *tessa* maximizes the correlation of the transcriptome and the TCR by deriving a position-wise weighting of the TCR features, which was uniform for all sequences in a dataset. *tessa* used the CDR3*β* sequence as the only input for representing the TCR. Therefore, we retrained *mvTCR* without the TCR*α* sequence to avoid an unfair advantage due to additional information. Except for clustering on donor 2, *mvTCR* showed better conservation of specificity on the 10x dataset. While we observed a large increase of 11.3% for kNN-prediction for donor 2, *mvTCR* only slightly surpassed the embedding derived from *tessa* by 2.45% for donor 1. On the Fischer dataset, *mvTCR* clustered significantly better for cell type (NMI, one-sided, paired t-test, p<1e-4) and reactivity (NMI, one-sided, paired t-test, p=0.026) with an increase in NMI of 25.4% and 8.1%, respectively. We attribute the gain in performance to *mvTCR’s* ability to integrate transcriptome information on a cell-level, while such information influences *tessa’s* embedding only on a dataset-level through its weighting factors. The conservation of characteristics captured in the gene expression makes *mvTCR* particularly suited for paired analysis of T cell repertoire data on transcriptomic- and TCR-level.

### *mvTCR* distinguishes activated from bystander T cells after SARS-CoV-2 infection

We used *mvTCR* to reanalyze Stephenson et al.’s dataset, which comprised more than 780,000 peripheral blood mononuclear cells (PBMCs) from 130 patients collected at three sites to study the coordinated immune response against SARS-CoV-2 infection [7]. From a total of 254,104 annotated T cells, 103,761 cells of 94 patients remained after filtering for complete TCRs (**Methods** Datasets). For each patient, the disease severity at the date of collection was provided as asymptomatic, mild (on ward, no oxygen required), moderate (on ward, oxygen required), severe (on intensive care unit (ICU), noninvasive ventilation), and critical (on ICU, intubation and ventilation). The study further contained a negative control group with cells from healthy donors, patients with other lung diseases, and donors with previously administered intravenous injections of lipopolysaccharide (LPS) to mimic inflammatory response. After training *mvTCR* on all T cells, we embedded the dataset into the shared representation (Fig. 2a). Separated groups of CD8^+^ effector T cells formed that expressed a high interferon (IFN) response score [18], which were captured by a fine-grained Leiden clustering in *mvTCR’s* representation (Fig. 2b,c). We observed that these Leiden clusters of cells with similar TCR and gene expression could not be identified in the transcriptomic space alone (Supplementary Fig. 4) demonstrating *mvTCR* synergistic embedding. After selecting clusters with highly significantly enriched IFN response scores (one-sided, unpaired t-test, p<0.001, Fig. 2d, Supplementary Data 2), we observed that all 15 resulting clusters contained almost exclusively (>99.5%) cells from donors with symptomatic SARS-CoV-2 infection withholding cells from asymptomatic infection or healthy, and negative control groups (Fig. 2e). Overall, the selected clusters consisted of cells from samples collected on average after 8.1 ± 4.0 days of symptom onset indicating an ongoing primary T cell response [19, 20]. Contrary, the cells of the remaining clusters originated from samples collected at a later date after symptom onset (11.8 ± 8.9 days). Generally, these clusters consisted of one or several expanded clonotypes with similar TCRs - 10 out of 15 clusters had significant lower inter-clonotype pairwise distance measured via *TCRdist* [21] (one-sided, unpaired t-test, p-values<0.05, Supplementary Data 2) - and similar phenotype - all clusters showed significant higher inter-cell correlation compared to remaining CD8^+^ effector cells (one-sided t-test, p-values<0.05, Supplementary Data 2). Since we observed similar phenotypes and functionality, we assumed that cells of a cluster were activated in the same fashion. While several clusters might be SARS-CoV-2 reactive, others might express a bystander response. Bystander T cells are activated by immune signaling without recognition of their cognate antigen [22] and were reported to play a crucial role in the different severity degrees of COVID-19 patients [23]. To assess T cell specificity, the TCRs were queried to *Immune Epitope Database (IEDB)* [24], which revealed 2,200 cells with possible cognate epitope matches (**Methods** Datasets). To reduce the number of false positives, we filtered out epitopes that were not predicted to bind to the corresponding donors’ HLA-types (Supplementary Data 3) via *MHCFlurry 2*.*0* [25] reducing the number of matches to 1,554 (Fig. 2f, Supplementary Data 4). Based on this query, we identified six SARS-CoV-2-specific and three bystander clusters. On average, the cells from SARS-CoV-2-specific clusters originated from earlier time points after symptom onset (7.7 days) compared to the bystander clusters (9.7 days) and the remaining clusters (11.7 days) again indicating an antigen-specific T-cell response (Fig. 2g).

**Figure 2.**
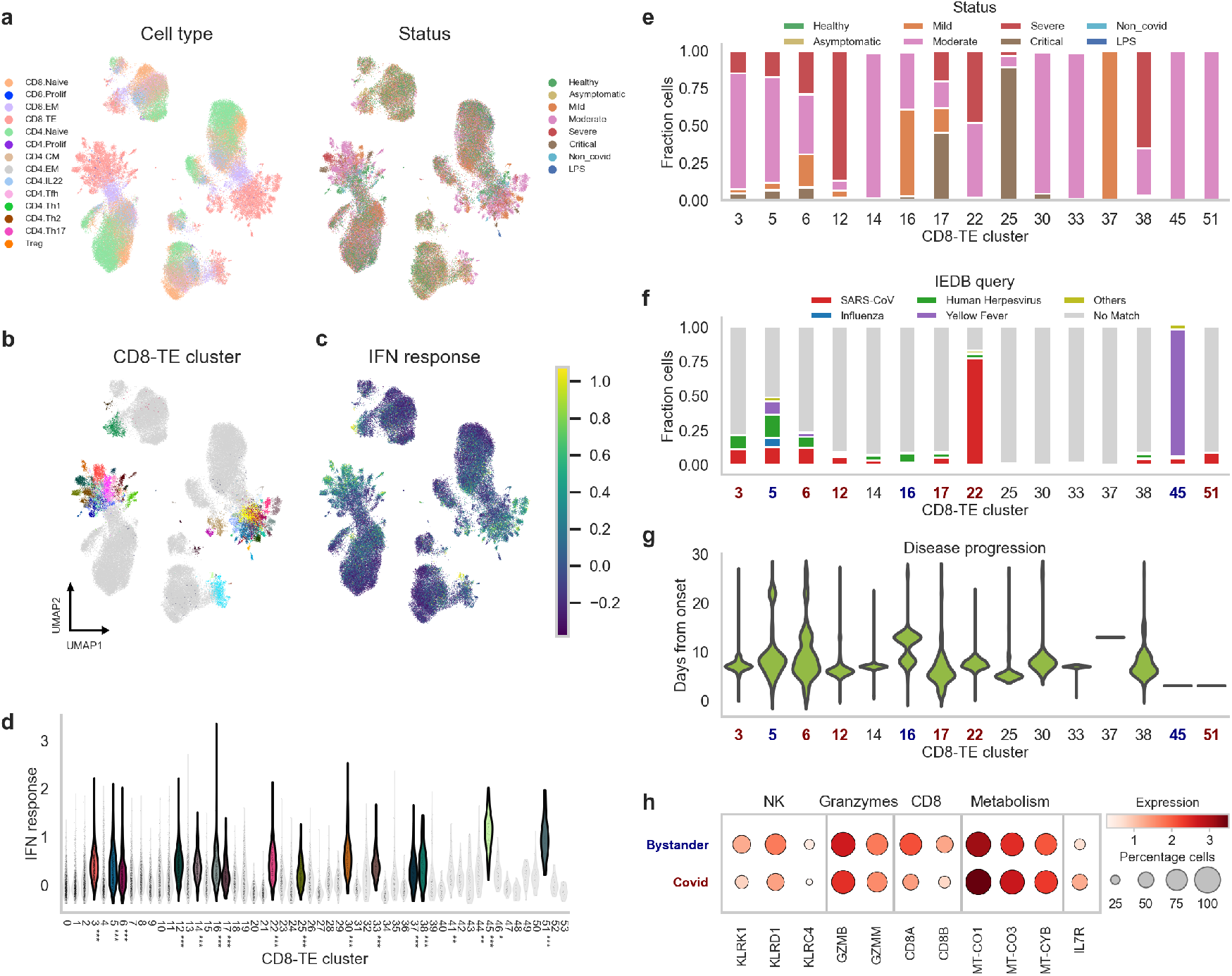
Joint embedding reveals hidden clusters in SARS-CoV-2 study. **a**, UMAP visualation of the joint embedding colored by annotation of cell type and status at day of hospital admission. **b** Effector CD8^+^ T cells form separating clusters. **c** UMAP colored by IFN response score, which is elevated in patients with symptomatic SARS-CoV-2 infection. **d**, Distribution of the IFN response score across the effector clusters with statistical significance (one-sided, unpaired t-test, *<0.05, **<0.01, ***<0.001). **e**, Patient status of cells from clusters with highly enriched IFN score. **f**, Specificity assignment of TCRs within the enriched clusters by query to the *IEDB* and predicted MHC restriction. **g**, Distribution of time after disease onset for the significant clusters. **h**, Selected differentially expressed genes between antigen-specific and bystander clusters.

Differential analysis between antigen-specific and bystander clusters (Fig. 2h, Supplementary Data 5) revealed several upregulated genes related to Natural Killer (NK) cells in the bystander clusters such as *KLRD1, NCR3*, and genes of the NK2G receptor group (*KLRK1* and *KLRC4*), which recognize stress-induced self-proteins [22]. Additionally, multiple granzymes (*GZMB, GZMM*, and *GZMK*) were upregulated indicating cytotoxic activity. This was coherent with previous studies on bystander activation in various diseases [22, 26], which linked elevated levels of *KLRK1* (*NKG2D*) and *NCR3* (*NKp30*) to an Interleukin 15 (*IL-15*) induced T-cell response in absence of TCR stimulation. Following *IL-15* exposure, CD8^+^ T cells adapt a NK-like phenotype and are able to kill targets in an innate-like fashion among others via cytotoxic granzymes. The antigen-specific clusters indicated downregulated CD8 expression (*CD8A, CD8B*) as previously reported for an active response of virus-specific CD8^+^ T cells eight days after infection [27]. Further, *IL7R* was expressed in 30.8% of the antigen-specific cells. While *IL7R* is downregulated in most CD8^+^ effector cells, it can be indicative of memory precursor effector cells (MPECs) which survive after viral clearing to form a long lasting immune memory [28]. Finally, several genes related to mitochondrial respiratory chain and oxidative phosphorylation (OXPHOS) (*MTCO1, MTCO2, MTCO3, MTCYB*) were observed in the antigen-specific cells. Even though there is a shift towards aerobic glycolysis after CD8^+^ activation, a parallel increase in OXPHOS level further contributes to the ATP production [29, 30], while both levels are further increased by a’peptide-MHC-induced activation [31].

In summary, *mvTRC* identified clusters showed compelling concordance of their activation pattern by TCR specificity, time after symptom onset, and differently expressed genes. These results demonstrate that *mvTCR* is able to identify striking clusters in T cell repertoires that would have been missed in unimodal analysis, and can, therefore, be applied to immunological relevant studies to uncover the interplay between T cell functionality and phenotype.

### Atlas level analysis is enabled through scArches-integration

Smaller-scale studies are now commonly integrated into large-scale atlases for knowledge transfer during annotation or analysis. Therefore, it is crucial for methods to scale to atlas-level datasets to leverage them as reference. To this extend, we applied *mvTCR* on a collection of 12 tumor-infiltrating lymphocyte (TIL) studies from different sources [32–43]. After filtering (**Methods** Datasets), the accumulated dataset contained a total of 722,461 T cells from 6 tissue sources covering 11 cancer types. During training of *mvTCR*, we excluded two studies to simulate query datasets [39, 40], while a reference was built from the remaining studies. To remove batch effects within the reference studies and query-atlas mapping, we extended *mvTCR* with architectural surgery (scArches) [14] (**Methods** Conditioning).

We trained *mvTCR* with and without scArches on the TIL dataset to evaluate the integration quality. Without scArches, the cells were embedded into fragmented clusters. While each cluster was mostly pure in regard to the same cancer type and origin (Supplementary Fig. 5), cells from the same biological labels separated into multiple clusters. In contrast, *mvTCR* with architectural surgery formed more contiguous clusters, while still preserving purity of cancer type and origin (Fig. 3a).

**Figure 3.**
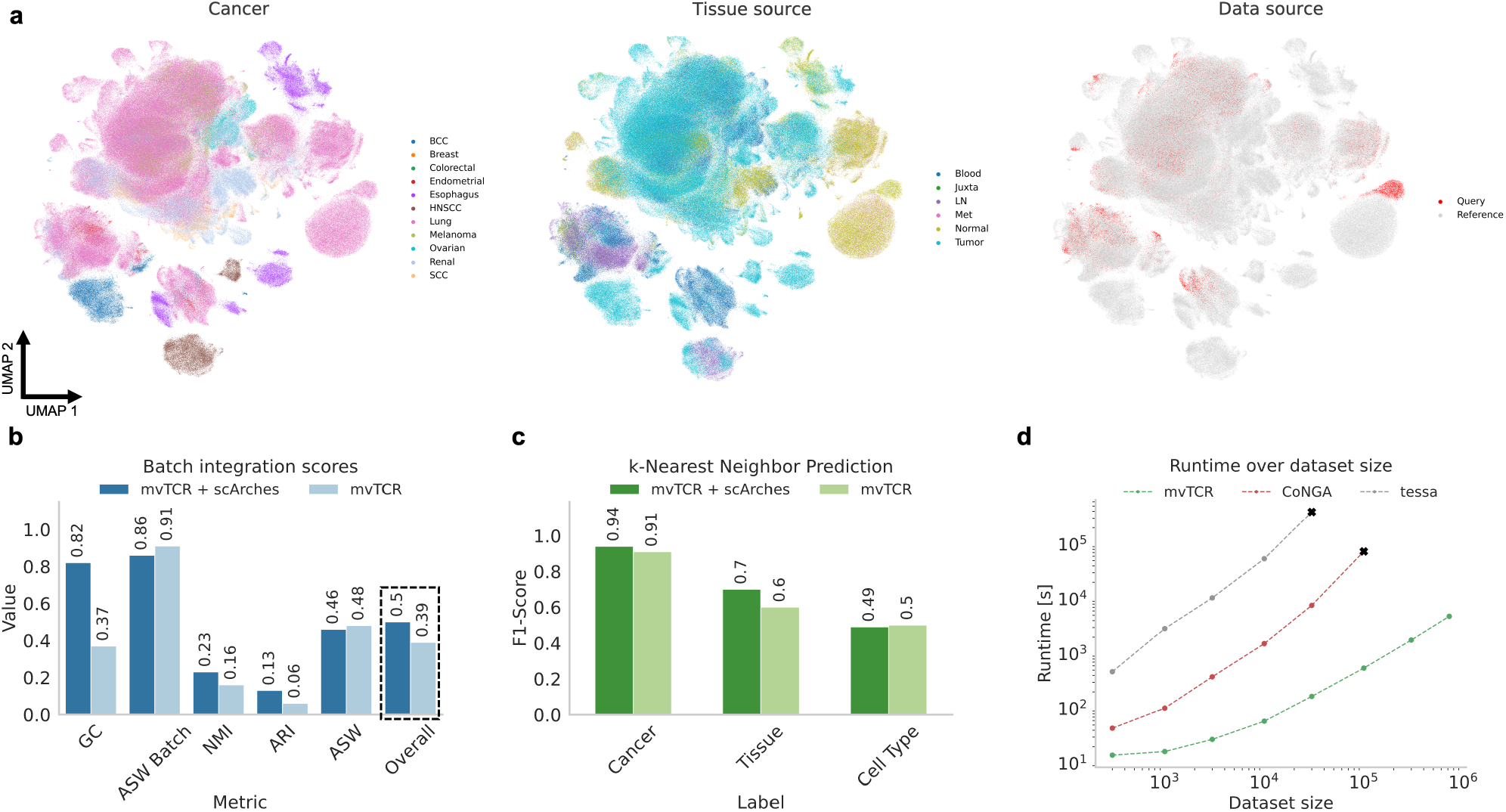
mvTCR enables query to atlas reference mapping. **a**, UMAP visualizations of the joint embedding colored by cancer tissue, tissue type, and query vs. reference for *mvTCR* with scArches. **b**, Comparison of data integration metrics [44] between *mvTCR* with and without architectural surgery (scArches). **c**, kNN prediction using the multimodal embedding as features to classify biological labels on the query dataset. **d**, Runtime in dependence to different dataset sizes, compared against *tessa* [9] and *CoNGA* [8].

The quality of data integration from different sources can be broken down into two dimensions: The correction of batch effects and the conservation of biological signals. These dimensions are interrelated as correction of batch effects often leads to removal of biological variation. To quantify batch correction effects, we use the Graph Connectivity (GC) score and Average Silhouette Width Batch (ASW Batch), while for estimating biological signal conservation we used Normalized Mutual Information (NMI), Adjusted Random Index (ARI), and Average Silhouette Width (ASW) on the cancer origin, origin of cells, and subtype of T cells (**Methods** Benchmarks). Using *mvTCR* with architectural surgery improved the averaged integration scores from 0.39 to 0.50 (Fig. 3b). This improvement was reflected in almost all individual metrics. The GC score improved drastically with architectural surgery, indicating better connectivity of subgraphs from cells with the same label. Furthermore, Leiden clusters had higher NMI values with cell annotations. Only the Average Silhouette Width dropped marginally, indicating a minor decrease in cluster compactness and distinctiveness for biological labels and less overlap between batches. Next, we assessed the quality of query-atlas mapping to annotate cells from held-out studies by a kNN-classifier predicting cancer type, tissue source, and cell type (Fig. 3c). Due to the reduction of technical noise, architectural surgery improved the predictions for cancer and tissue source, while being almost on par for cell type annotations. Therefore, we conclude, that *mvTCR* can efficiently remove batch effects between query and atlas sets, while preserving the biological signal between studies.

To compare the scalability of *mvTCR* with other established methods integrating gene expression and TCR information, we assessed the execution time of *mvTCR, CoNGA* [8], and *tessa* [9] on subsampled dataset sizes (**Methods** Benchmarks). For all dataset sizes, *mvTCR* was significantly faster than *CoNGA* and *tessa* (Fig. 3d). In comparison to *CoNGA, mvTCR* was up to 135 times faster (N=100,000 cells - *mvTCR*: 545 s, *CoNGA*: 73,516 s), while *tessa* needed up to 2,353-fold more time (N=30,000 cells - *mvTCR*: 166 s, *tessa*: 389,578 s). Besides runtime, the memory requirements for *CoNGA* and *tessa* were also higher due to pairwise comparisons. On the specified machine used for runtime benchmarking, both *CoNGA* and *tessa* exceeded the memory available (256 GB) for dataset sizes of 30,000 and 100,000 cells, respectively, demonstrating *mvTCR’s* great scalability to atlas-scale dataset integration.

## Discussion

As more and more paired single-cell TCR- and RNA-seq datasets are being collected, scalable methods are needed to leverage both modalities for joint analyses. Here, we presented *mvTCR* - a multiview Variational Autoencoder - to enable large-scale integration of paired TCR and transcriptome data for single-cell studies of T cell repertoires. We showed that *mvTCR* can preserve both cell state and function by incorporating both modalities into its shared representation while capturing more information than unimodal representations as demonstrated in antigen and avidity prediction, reference mapping, and clustering. Furthermore, we demonstrated *mvTCR*’s scalalability to atlaslevel datasets for T cell reference construction. Especially when combined with scArches, *mvTCR* is capable of correctly mapping new multimodal studies into reference atlases enabling systematic extensions of references and automated analysis of query datasets.

*mvTCR* has robust performance on multimodal T-cell datasets. Yet, it is limited to paired data, where gene expression is available in combination with *α*- and *β*-CDR3 of the TCR. A natural extension would be to incorporate other modalities such as chromatin accessibility and surface protein abundance as recently proposed by non-TCR-aware single-cell multimodal integration methods [45– 47]. Adding a supervised component to the model to predict epitope specificity could further guide the network’s training and improve the multimodal representation. Additionally, interpretability methods could be applied to detect TCR and transcriptome characteristics indicative of the cell’s functional role. Furthermore, pre-trained TCR language models [48, 49] in combination with VDJ-gene usage encoding could be used to improve the TCR representation [50, 51]. Lastly, *mvTCR* can be extended to joint B-cell receptor data, though the modeling of somatic hypermutations needs to be carefully considered to accurately represent B-cell lineages. As a technical limitation, adjusting the contribution of each modality requires repeated training of *mvTCR*, ideally by an additional hyperparameter search. Even though an automated selection of network parameters could partially prevent retraining, the desired contribution of each modality is dependent on the study-specific analysis objective.

In conclusion, we presented *mvTCR* as a model for analysing T cell repertoires in the context of infectious disease, tumors, and therapies. We envision that the integration of TCR and transcriptome via *mvTCR* will uncover hidden interdependencies between the two modalities and identify functional related T-cell sub-clusters, that would remain hidden in separate analysis of the T-cell response, thereby contributing to our understanding of T-cell modulation in healthy and disease.

## Code availability

The software code including tutorials is available at https://github.com/SchubertLab/mvTCR. The code to reproduce the results of this manuscript can be accessed under https://github.com/SchubertLab/mvTCR_reproducibility. All trained models used for this manuscript can be downloaded from Zenodo via https://doi.org/10.5281/zenodo.7215447.

## Data availability

All datasets used in this paper are publicly available. The 10x dataset (https://www.10xgenomics.com/resources/datasets?query=&menu[products.name]=Single%20Cell%20Immune%20Profiling, Section *Application Note - A New Way of Exploring Immunity*, accessed March, 7^th^, 2021), and the SARS-CoV-2 dataset (https://www.covid19cellatlas.org/index.patient.html, Section *COVID-19 PBMC Ncl-Cambridge-UCL*, accessed February, 2^nd^, 2022) were downloaded from the linked repositories as described in their original publication. The Fischer dataset can be accessed via NCBI GEO under the accession number *GSE171037*. The samples contained in Borchdering dataset stem from a collection of studies. A processed version of this data was downloaded as described in https://github.com/ncborcherding/utility/tree/dev from TIL (accessed December, 20^th^, 2021).

## Author Contributions

M.L. and B.S. conceived the project. F.D. and Y.A. performed research, implemented models, and performed analysis. L.D. and R.L. tested the models and assisted with the analysis. M.H., S.T, and F.T provided feedback on the method and the manuscript. M.L. and B.S. supervised the research. All authors wrote the manuscript.

## Acknowledgments

This work was supported by the BMBF grant DeepTCR (#031L0290A), by the Helmholtz Association’s Initiative and Networking Fund on the HAICORE@FZJ partition, and Helmholtz International Lab “Causal Cell Dynamics”. Y.A. and F.D. are supported by the Helmholtz Association under the joint research school “Munich School for Data Science - MUDS”. M.L. acknowledges financial support from the Joachim Herz Stiftung. L.M.D is supported by European Union’s Horizon 2020 research and innovation programme under the Marie Skłodowska-Curie grant agreement No 955321.

## Competing interests

F.J.T. reports receiving consulting fees from Roche Diagnostics GmbH and Cellarity Inc., and ownership interest in Cellarity, Inc. Y.A. acknowledges financial support by JURA Bio, Inc.

## Methods

### mvTCR

*mvTCR* was trained on paired single-cell TCR sequences and RNA-seq datasets. A dataset 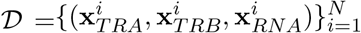 consists of 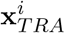 and 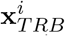 representing the *α*- and *β*-chain of the TCR and 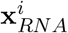 indicating the expression for each cell *i*. 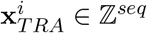 and 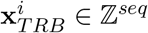 contain the amino acid sequence of the highly variable CDR3. Both sequences are tokenized and zero-padded to the maximal sequence length *seq* present in 𝒟. In the following, 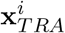 and 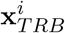 are summarized as 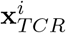 when both chains are considered. 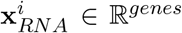 comprises the 5,000 most highly variable gene, whose read counts were normalized and log1p-transformed.

*mvTCR* encodes the TCR sequences 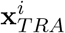 and 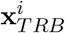 via the two encoder *E*_*TRA*_ and *E*_*TRB*_, respectively, to obtain the lower-dimensional representations:

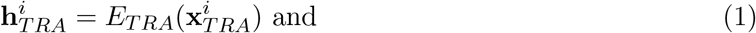

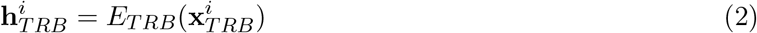

of size *h*/2. Both representations are then concatenated to form the TCR embedding 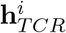. Similarly, 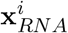 is transformed via the encoder *E*_*RNA*_ to the embedding:

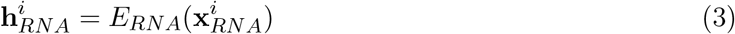

of size *h*. Next, both embeddings are combined via different versions of the mixture model *M* leading to the shared latent distribution:

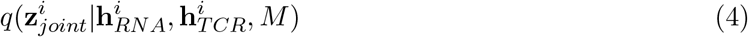

of size *h*. All downstream analysis and benchmark tests were performed on 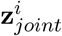. The networks *D*_*RNA*_, *D*_*TRA*_, and *D*_*TRB*_ decode the embeddings to the reconstructions:

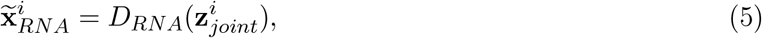

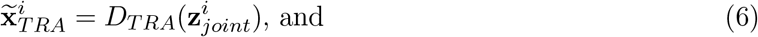

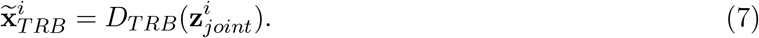

### Network Structure

*mvTCR* consists of several networks, specifically, the encoders and decoders for TCR and transcriptome, and different variants of the mixture module for fusing both modalities.

#### RNA networks

Following [13, 14], *E*_*RNA*_ uses the architecture of a multi-layer perceptron. Each layer was build by a block containing a fully connected layer, followed by batch-normalization [52], leaky ReLU activation, and a dropout layer [53]. Via a linear layer, the output was transformed to *h. D*_*RNA*_ similarly consisted of these blocks with a final layer with linear activation function obtaining the original input size of 5,000 genes.

#### TCR networks

Based on its performance on sequence data in Natural Language Processing, we employed the transformer architecture [54] for extracting features from 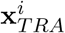 and 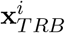 via the encoders *E*_*TRA*_ and *E*_*TRB*_. The output of each encoder was transformed, separately, via a fully connected layer with linear activation function to *h/*2. While *E*_*TRA*_ and *E*_*TRB*_ followed the same architecture, they did not share their weights, to allow each network to focus on unique features of their respective input. The shared representation 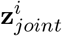 was up-sampled via a fully connected linear layer, before yet again transformer blocks were used for the decoding networks *D*_*TRA*_ and *D*_*TRB*_. Finally, a linear layer with softmax activation function reconstructed the amino acid sequence.

For fusing the two modalities, three different versions of the mixture models *M* were implemented. Additional, models trained on either the transcriptome or the TCR modality were used as a unimodal baseline model.

#### Concatenation

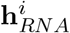 and 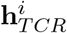 are concatenated to a representation of size *h* ∗ 2, which is passed to an additional shared encoding network *E*_*joint*_. This network consists of the same blocks described above and estimates the mean ***µ*** and standard deviation ***σ*** of the normal distribution 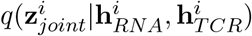 from which ***z***_*joint*_ is sampled via the reparameterization trick [55]. Note, that ***µ*** is used for all downstream analysis throughout this paper.

#### Product of Experts (PoE)

Contrary to the concatenation model, PoE uses additional encoder networks *E*_1_ and *E*_2_ to obtain mean (***µ***_1_ and ***µ***_2_) and standard deviations (***σ***_1_ and ***σ***_2_) for each modality, individually, resulting in the latent distributions 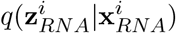 and 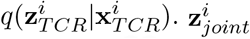 is sampled via the reparameterization trick from the product of these distributions. ***µ*** and ***σ*** can be calculated from its closed form solution

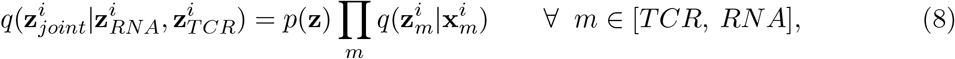

where *p*(*Z*) is an univariant Gaussian prior with zero-mean [15]. To motivate the linkage of knowledge between both modalities, the reconstruction was calculated from the shared as well as the modality-specific latent distribution by the same decoder.

#### Mixture of Experts (MoE)

As the PoE, MoE calculates individual latent distribution, which are then both used to reconstruct each modality. This forces the encoder networks to have similar predictions for TCR and transcriptomic input [16]. For downstream analysis, the average of both distributions

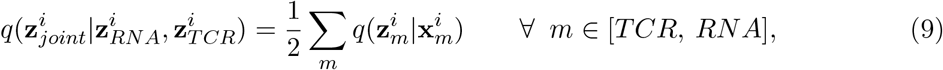

is used. If not stated otherwise, this mixture module was used throughout this paper.

#### Unimodal models

An encoding network of fully connected blocks described above estimated the mean ***µ*** and standard deviation ***σ*** of the normal distribution 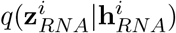 or 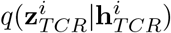 from 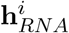 or 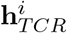, respectively. The reconstruction was calculated from the sampled 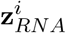 or 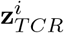.

### Conditioning

To integrate query datasets into a trained reference atlas model, we followed a similar approach as Lotfollahi et al. [14]. First, the model is trained on the samples from the atlas datasets to build a reference model. Since the reference atlas may consist of multiple different studies, batch effects can occur between those. To counter batch effects, *mvTCR* is conditioned towards the studies. Since the MoE mixture module is used in our experiments for query to reference atlas mapping, we describe the procedure for this version of the mixture model only. Let **c**_*j,atlas*_ be a trainable embedding of dimensionality *D*_*c*_ for each atlas study *j* representing the difference between studies. For each cell *i* the corresponding conditional embedding is concatenated to the hidden representations 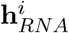 and 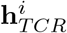 before calculating the individual latent distributions. Similarly, the same embedding **c**_*j,atlas*_ is concatenated towards the individual latent representations 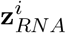 and 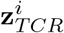 before passing them into the corresponding decoders. After the training converged for the reference dataset, the query is integrated using architectural surgery [14]. All parameters of the reference model are frozen and embeddings **c**_*j,query*_ for the new query studies are randomly initialized. Only these embeddings **c**_*j,query*_ are trained on the new query datasets. This procedure reduces the number of parameters to be trained by multiple orders of magnitudes while preventing catastrophic forgetting.

### Training

The models are trained on the weighted sum of reconstruction losses encouraging the conservation of the input data and regularization losses, which shaped the properties of the latent distribution.

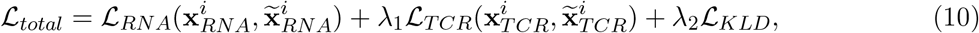

where

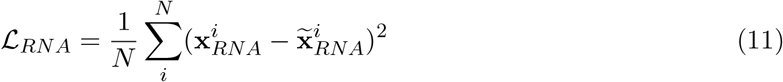

is the mean squared error, and

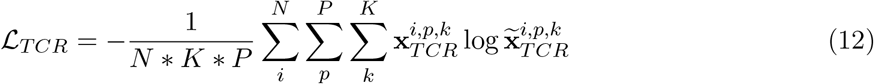

the Cross Entropy loss over the sequence encodings for each cell *i* over each amino acid label *k* per position *P* in the TCR. The Kullback-Leibler divergence loss

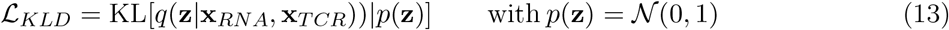

constrains the latent distribution to resemble a univariant, zero-mean Normal distribution and is applied to all latent distributions of the respective mixture model. The loss is minimized by the ADAM optimizer with the learning rate as a hyperparameter [56]. The datasets is split into different subsets before training the model. The loss function ℒ_*total*_ is reduced by optimizing all subnetworks of the model jointly on the training data until the validation loss stops decreasing for 5 epochs or a maximum of 200 epochs is reached. Since datasets often contained highly expanded clonotypes, the TCR encoder and decoder focuses on over-represented sequences. Therefore, we oversample cells with low-frequency TCRs in the training set of the joint and TCR models by sampling with a probability

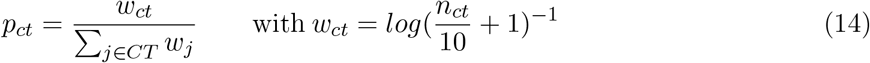

for each clonotype *ct* from the set of all clonotypes *CT*.

For benchmarks (Fig. 1c-f, Supplementary Fig. 3), an additional test set of 20% was used to evaluate the performance on unobserved data. Training, validation, and test sets were constructed randomly on a clonotype-level, i.e., cells with the same TCR input sequence were exclusively contained in a single subset.

For evaluating atlas-level integration (Fig. 3a-c), two studies [39, 40] containing cells from lung cancer patients were held out of the accumulated dataset and 20% of the remaining data was used as validation set.

To compare the running times with other multimodal methods (Fig. 3d), *mvTCR* was trained on random subsets. Again 20% of data was used as validation set to measure the time for evaluation calculations. Since the model converged after 20 epochs on the full dataset, this number was held constant over all subsets, i.e., no early stopping was performed.

### Hyperparameter Optimization

To select the best model structure, we perform optimization of all hyperparameter of the architecture via *Optuna* [57]. Depending on the information available, different performance metrics are optimized to obtain the best model over different training runs. When cell-level epitope specificity information is available (10x dataset), the models are evaluated by their ability to capture specificity in the embedding. Specifically, the weighted F1-Score for predicting epitope specificity via a kNN classifier (k=5) is evaluated between training (atlas) and validation (query) set. For the remaining datasets, the models are optimized on how well they preserve cell type and clonotype measured by consistency of their label in the embedding within the validation set. Here, the weighted F1-Score is calculated for predicting both annotations by its nearest neighbor. Finally, the models are optimized towards maximizing a weighted sum of clonotype and cell type preservation, which further enables us to determine the dataset-specific degree to which the shared embedding is influenced by each modality. Models on all datasets were optimized for 48 GPU-hours, except on the TIL dataset, where the training time was increased to 96 GPU-hours due to the dataset size. All results, were obtained from the best performing model on these performance metrics.

### Avidity Prediction

A prediction head is fitted to predict the pMHC tetramer read counts **a**^*i*^ of the most abundant eight epitopes in the 10x dataset. This additional neural network consists of the fully connected blocks with an exponential activation layer. Using the mean squared logarithmic error

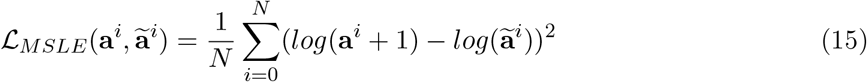

between the ground truth and the predicted avidity **a**^*i*^ the models are trained with ADAM optimizer and early stopping (patience of 10). The hyperparameters are optimized by *Optuna* on 100 training runs.

### Benchmarks

We compare the different multi- and unimodal models of *mvTCR* and *tessa* with the following metrics:

#### F1-Score

the performance for predicting cell-level labels with a k-nearest neighbour classifier is evaluated by the harmonic mean between precision and recall. To aggregate performance over all labels *L*, the F1-Score

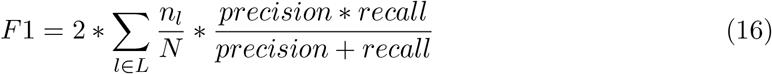

is weighted by the class support. This metric is applied for predicting antigen specificity on the 10x dataset and for cell type, tissue source, and tissue on the TIL dataset.

#### Normalized Mutual Information (NMI)

The NMI is used to compare the overlap between clusters *C* in the shared embedding and cell labels *L* via

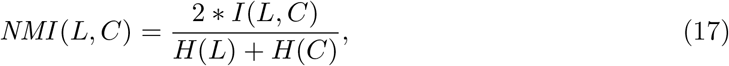

which normalizes the mutual information *I*(*L, C*) by the entropies *H*(*L*) and *H*(*C*). To derive clusters in the latent space, Leiden clustering is applied for different resolution factors (0.01, 0.1, 1.0) and the maximal NMI value between labels and annotation is reported. The NMI is reported for evaluating clustering of antigen specificity in the 10x dataset, cell type, and reactive clonotypes in the Fischer dataset. On the TIL dataset the cancer type, tissue source, and the cell type is used as labels. The best performing resolution out of (0.01, 0.03, 0.1, 0.3, 1.0, 3.0) is used for each label individually.

#### Mean Squared Logarithmic Error (MSLE)

Following Fischer et al. [58], the MSLE as described in Eq. 15 is used to evaluate the prediction of avidity counts in the 10x dataset.

#### R^2^-Score

The Coefficient of determination (R^2^-Score) between log1p-transformed network output and avidity counts is used as an additional metric for evaluating predicting binding strength on the 10x dataset.

#### Graph Connectivity Score

The graph connectivity score quantifies how well cells of the same biological label *l* ∈ *L* are connected in the kNN graph on the embedding space. Following Luecken et al. [59], this metric is calculated as

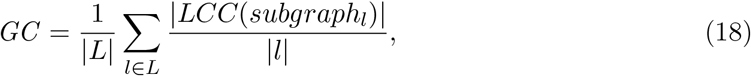

where *LCC*(*subgraph*_*l*_) indicates all cells within the largest connected component of type *l*, |*l*| the number of cells from type *l* and |*L*| the number of labels. The average over all labels are taken and the metric ranges from 0 to 1. A score of 1 indicates that all cell of type *c* are connected within one kNN graph.

#### Adjusted Random Index

The ARI compares the overlap of predicted clusters and biological labels. It assesses both correct overlaps and simultaneously counting correct disagreements. Similar to the NMI score, we use Leiden clustering with the following resolutions (0.01, 0.03, 0.1, 0.3, 1.0, 3.0) and retain the resolution with the best ARI on the labels - cancer type, tissue source, and the cell type.

#### Average Silhouette Width

This score measures the average distance in embedding space of one cell to all other cells, while distinguishing between cells of the same and different types. Following Luecken et al. [59] the score is normalized to range between 0 and 1, where 1 indicates that cells are well clustered within each type and separated from clusters of other types. Biological signals should be conserved after integration, hence, a score of 1 is desired. Again, on the TIL dataset the cancer type, tissue source, and the cell type is used as labels. On the other hand, for batch effect correction, individual studies should be as indistinguishable as the biological variation allows. Therefore, Luecken et al. modified the calculation, so that 1 represents a perfect overlap of batches [59]. In the experiment, each data source is treated as a unique batch.

Benchmarking tests are performed under the following settings:

#### 10x dataset

*Optuna* optimized the hyperparameters for predicting kNN-prediction of antigen specificity. The model is retrained and evaluated on dataset splits on five different seeds during the benchmark tests (Fig. 1c, Supplementary Fig. 3a) to enable statistical testing. The avidity prediction (Fig. 1d, Supplementary Fig. 3b) is conducted on the same training, validation, and testing splits as indicated above.

#### Fischer dataset

as for the 10x dataset, the models are trained five times on different dataset splits (Fig. 1e). The hyperparameter are adapted via *Optuna* to preserve cell type and clonotype at a ratio of one-to-one.

#### Comparison to *tessa*

We evaluate the performance of *mvTCR* against *tessa* as a baseline model (Fig. 1f, Supplementary Fig. 3c). However, *tessa* takes the CDR3*β* sequence as sole input of the TCR. Therefore, we retrain *mvTCR* without the CDR3*α* sequence for the 10x and Fischer dataset to avoid the advantage of additional information. Here, we directly use 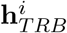 as 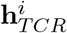 instead of concatenating it with 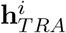. In this setting, we consider clonotypes as cells with identical CDR3*β* sequence to avoid the same TCR information in the different subsets of the data. The remaining training follows the description above. After applying the *tessa* algorithm, kNN predictions are evaluated on the resulting weighted TCR embedding. The cluster annotation provided by *tessa* is evaluated using the NMI-based metrics.

#### Query to reference mapping

Since the MoE model worked best, we compared this variant with and without architectural surgery [14] on the TIL dataset. Both are optimized using *Optuna* to determine the best hyperparameter sets to preserve cell type and clonotype with a ratio of 10 to 1.

#### Runtime vs dataset size

In this experiment, we compare the runtime of *mvTCR* with two concurrent methods integrating gene expression and TCR information - *tessa* [9] and *CoNGA* [8] on a computer with 2x Intel Xeon Gold 6226R (in total 32 Cores), 256 GB RAM, and 1 Nvidia Tesla V100. The same subsets of the full TIL dataset are used for all experiments. We first determined the number of training epochs needed for *mvTCR* to converge on the full dataset and keep this number (20 epochs) constant over all subsets and no early stopping was performed. The runtime corresponds to the training time for 20 epochs without counting the preprocessing and inference time. Similarly for *tessa* and *CoNGA*, we also excluded the preprocessing time. The time for *tessa* is counted from running the BriseisEncoder and *tessa* clustering. *CoNGA* is run as provided in their example script and the runtime as defined by the original authors is logged.

### Datasets

#### 10x dataset

The dataset for all four donors was downloaded from [10x Genomics](https://www.10xgenomics.com/resources/datasets?query=&page=1&configure%5Bfacets%5D%5B0%5D=chemistryVersionAndThroughput&configure%5Bfacets%5D%5B1%5D=pipeline.version&configure%5BhitsPerPage%5D=500&menu%5Bproducts.name%5D=Single%20Cell%20Immune%20Profiling) under the section *Application Note - A New Way of Exploring Immunity*. Following [44], we performed quality control using *Scanpy* [60], which can shortly be described as following: to remove lysed and dying cells, we filtered cells exceeding a fraction of 20% mitochondrial reads. Additionally, we only considered cells within the span of 1,000-10,000 reads counts with a minimum of 500 genes per cell. Genes reported for less then 10 cells were removed from the dataset. Doublets were filtered using *Scrublet* at a threshold of 0.05 [61]. The gene expression data was normalized to 10,000 reads per cell, followed by log1p-transformation and the reduction to the 5,000 most highly variable genes. Additionally, the specificity annotation suggested in the publication note was added. All cells not expressing a full TCR consisting of one *α*- and *β*-chain were removed from the datasets since *mvTCR* requires paired information. To ensure correct matching between TCR and specificity in our benchmark data, we further removed all cells expressing multiple TCRs. A clonotype ID was assigned grouping cells with identical *α*- and *β*-chain. For better quantification during the benchmark studies, this dataset was reduced to T cells with reported binding to the most abundant eight antigens excluding non-binders. The cytotoxicity score (Fig. 1b, Supplementary Fig. 1 and 2), was calculated by the mean of the normalized marker genes described in [18].

#### Fischer dataset

The filtered, normalized, and log1p-transformed dataset of Fischer et al. [6] was downloaded from [NCBI GEO](https://www.ncbi.nlm.nih.gov/geo/query/acc.cgi?acc=GSE171037). Cells with missing *α*- or *β*-chain were removed from the dataset. Clonotypes were assigned for the remaining cells. Finally, we selected the 5,000 most variable genes for training *mvTCR*.

#### SARS-CoV-2 dataset

We obtained the SARS-CoV-2 dataset from the Covid19 Cell Atlas under the section *COVID-19 PBMC Ncl-Cambridge-UCL* and manually joined transcriptomic data with TCR information. Quality control, normalization, and log1p-transformation was already performed in the original publication. We reduced the dataset to the 5,000 most variable genes, filtered for incomplete TCRs, and assigned the clonotype. The models were selected to equally preserve cell and clonotype. The database query of TCRs to the *IEDB* (Fig. 2d) was performed via *Scirpy* version 0.10.1 [62] using the Levenshtein distance with threshold 1. For predicting HLA binding affinity to the resulting epitopes, we used *MHCFlurry 2*.*0* with a threshold of 500 nM [25]. The IFN response score was calculated as the mean of the normalized marker genes described in Szabo et al. [18]. CD8^+^ effector T cell clusters with elevated IFN response were assigned as antigen-specific, if more than 5% of the cluster’s cells and more than 50% of the epitope matches of the cluster stemmed from SARS-CoV-2 variants. Similarly, bystander clusters were defined as exceeding these thresholds with non-Covid related epitopes. Differential gene expression was calculated between antigen-specific and bystander clusters via a t-test with Benjamini-Hochberg correction using scanpy. Only genes with an adjusted p-value smaller 5% and a log-fold change greater 0.25 were reported.

#### Tumor-Infiltrating Lymphocyte dataset

The Tumor-Infiltrating Lymphocyte (TIL) dataset consisted of a collection of studies, which were downloaded as described under https://github.com/ncborcherding/utility/tree/dev. Transcriptome and TCRs were manually merged and cells without complete TCR were filtered. Additional, annotated doublets and genes present in less than 100 cells were removed. The data was normalized to 10,000 counts per cell, log1p-transformed, reduced to 5,000 most variable genes, and annotated with clonotype. After filtering, the dataset contained 722,461 T cells.

## Supplementary Figures

**Supplementary Figure 1.**
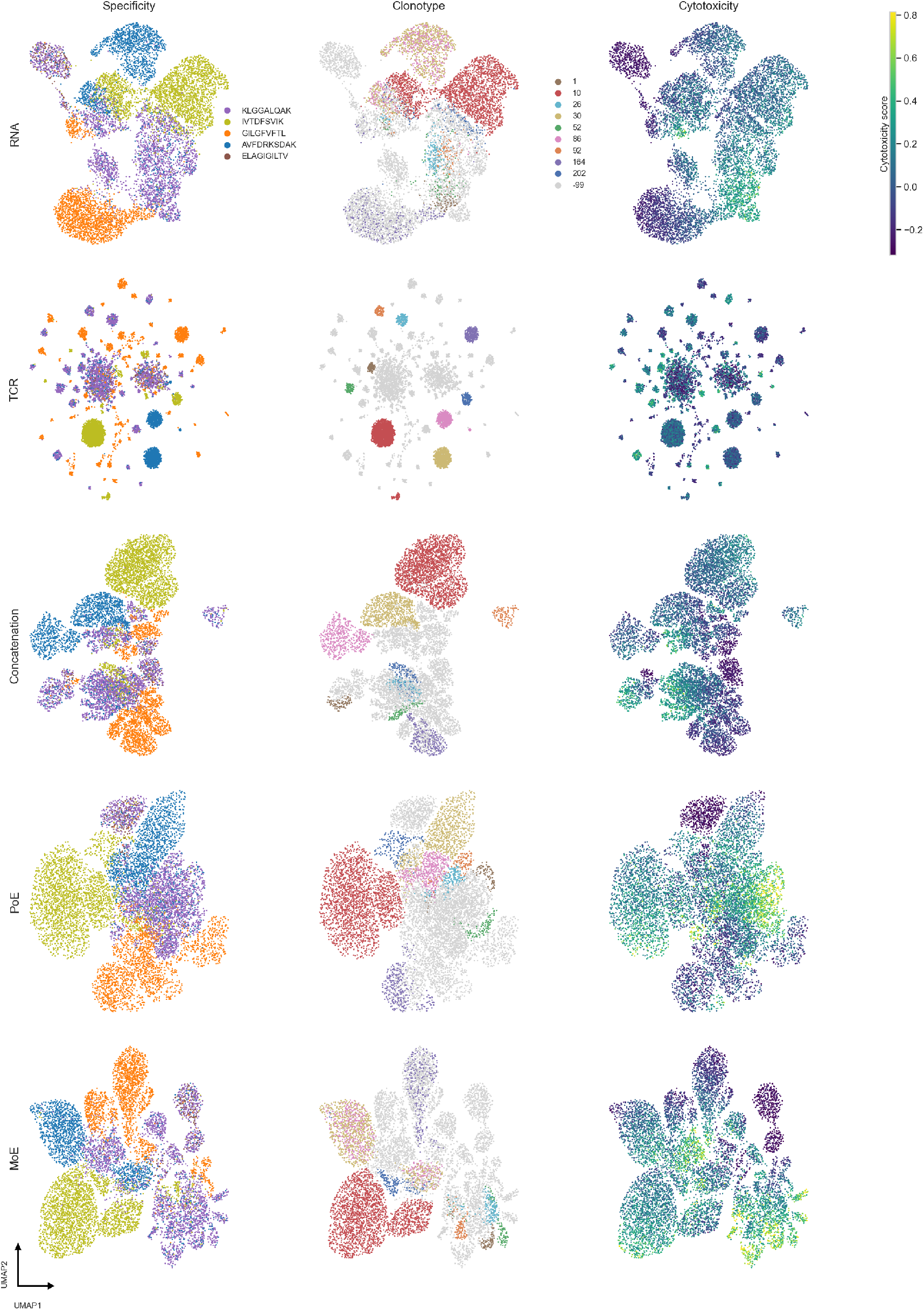
Embeddings for 10x Genomics donor 1. UMAP visualisations of the embeddings from different uni- and multimodal models colored by epitope specificity, ten largest clonotypes, and a cytotoxicity score as an example of cell state.

**Supplementary Figure 2.**
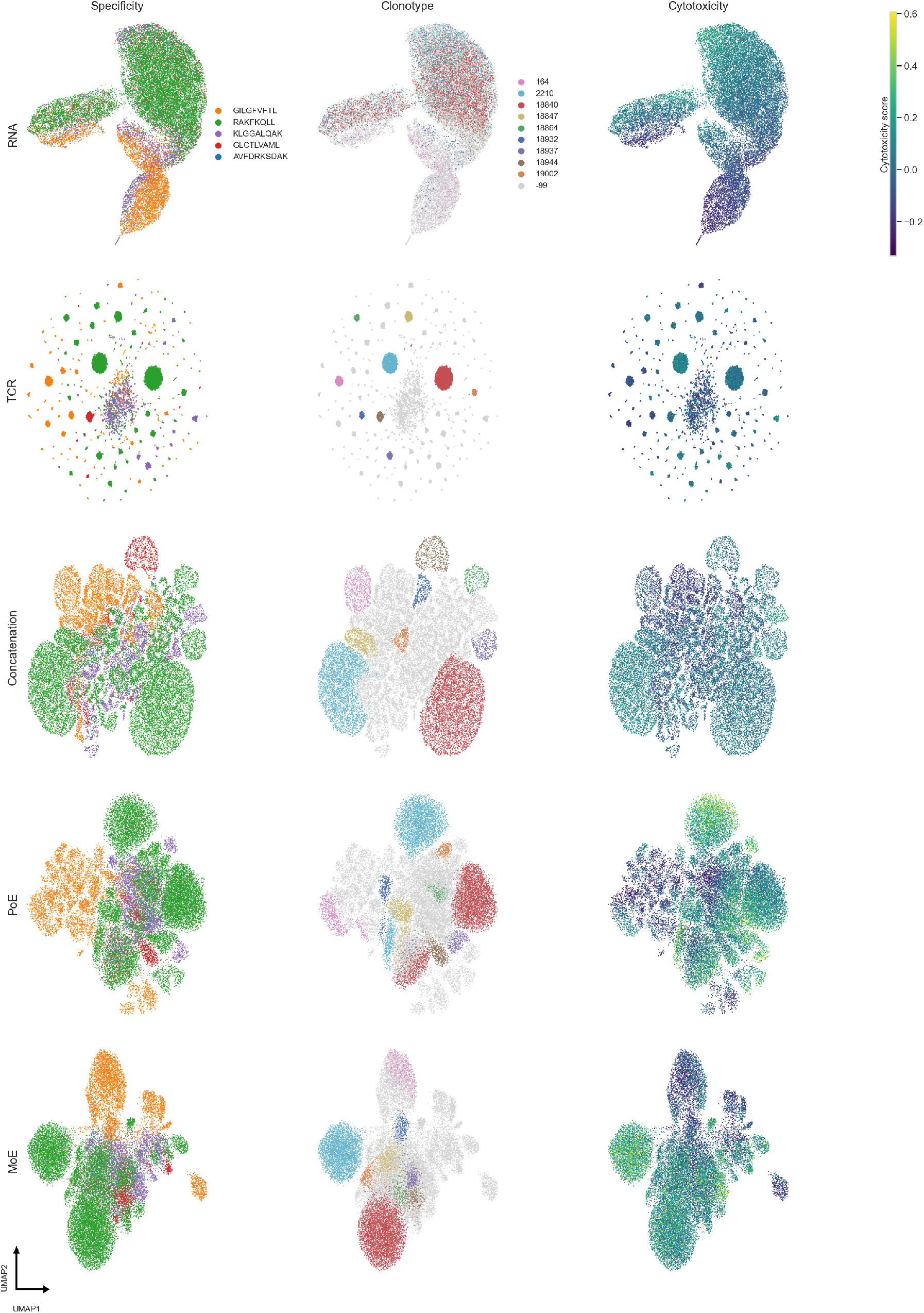
Embeddings for 10x Genomics donor 2. UMAP visualisations of the embeddings from different uni- and multi-modal models colored by epitope specificity, ten largest clonotypes, and a cytotoxicity score as an example of cell state.

**Supplementary Figure 3.**
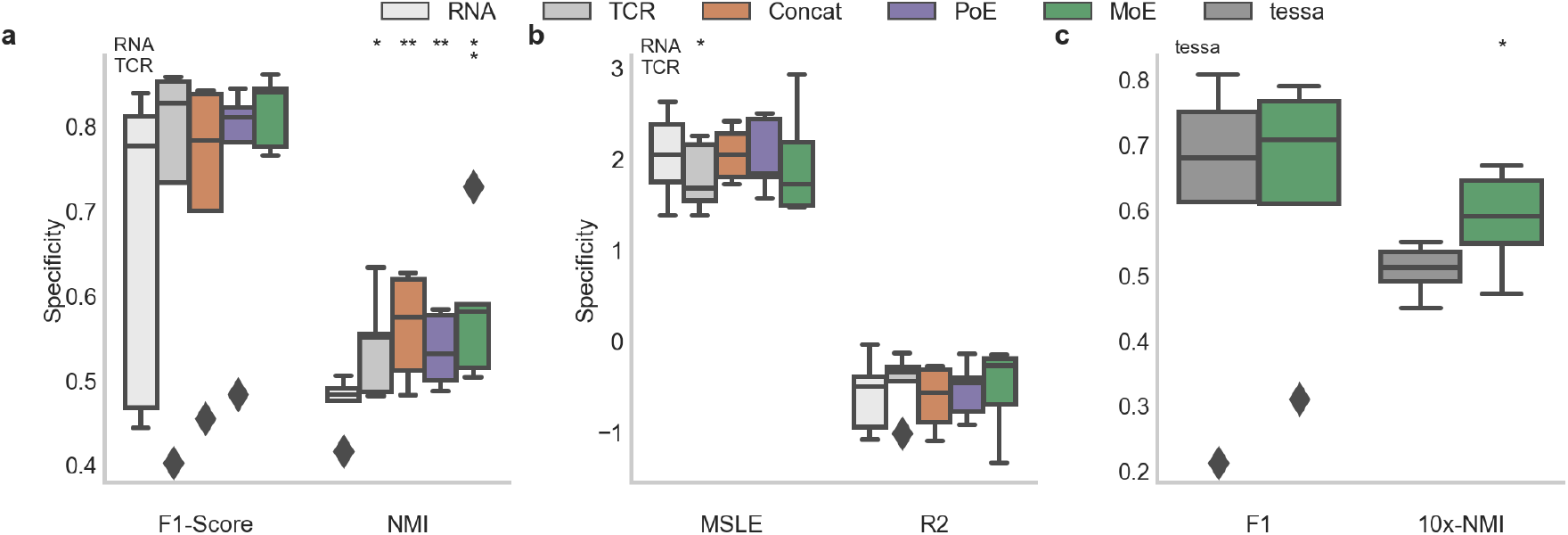
Quantitative evaluation for 10x Genomics donor 1. Statistical significance to the corresponding uni-modal embedding is calculated via one-sided, paired t-test (p-values: *<0.5, **<0.01, ***<0.001). The boxplot represents the quartiles and median line, while the whiskers extend to the full value range excluding outliers. **a**, Comparison of uni- and multi-modal models for capturing of specificity by atlas-query prediction (weighted F1-Score) and clustering (Normalized Mutual Information). **b**, Comparison of uni- and multi-modal models for avidity prediction measured by mean squared logarithmic error and R^2^ value. **c**, Comparison between MoE embedding trained only on gene expression and CDR3*β* and *tessa* on the tasks defined in b.

**Supplementary Figure 4.**
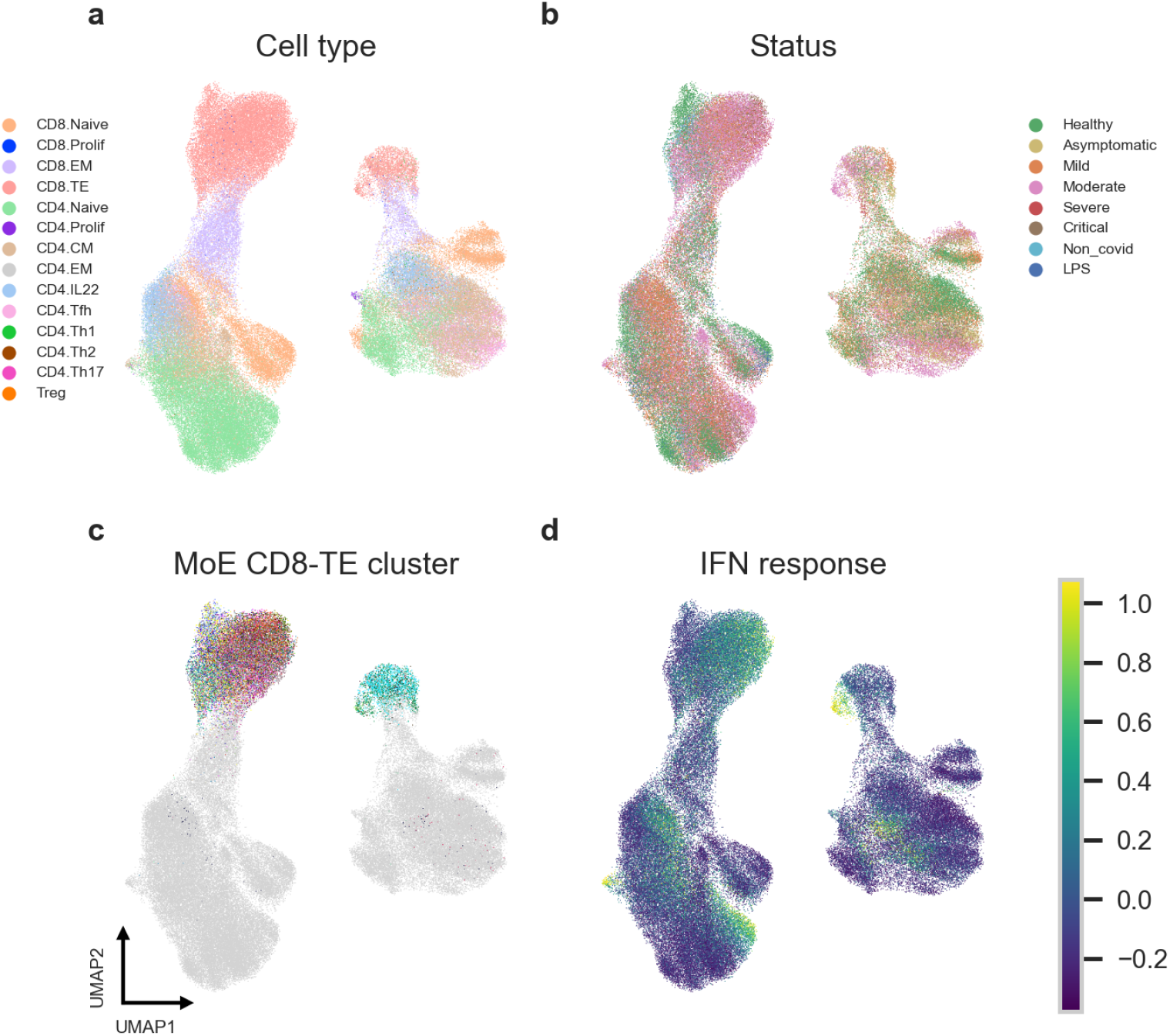
Embeddings of the RNA-model on the SARS-CoV-2 dataset. UMAP visualation colored by cell type (**a**), patient status (**b**), clusters of CD8^+^ T cells defined on the MoE-embedding (**c**), and IFN response score (**d**).

**Supplementary Figure 5.**
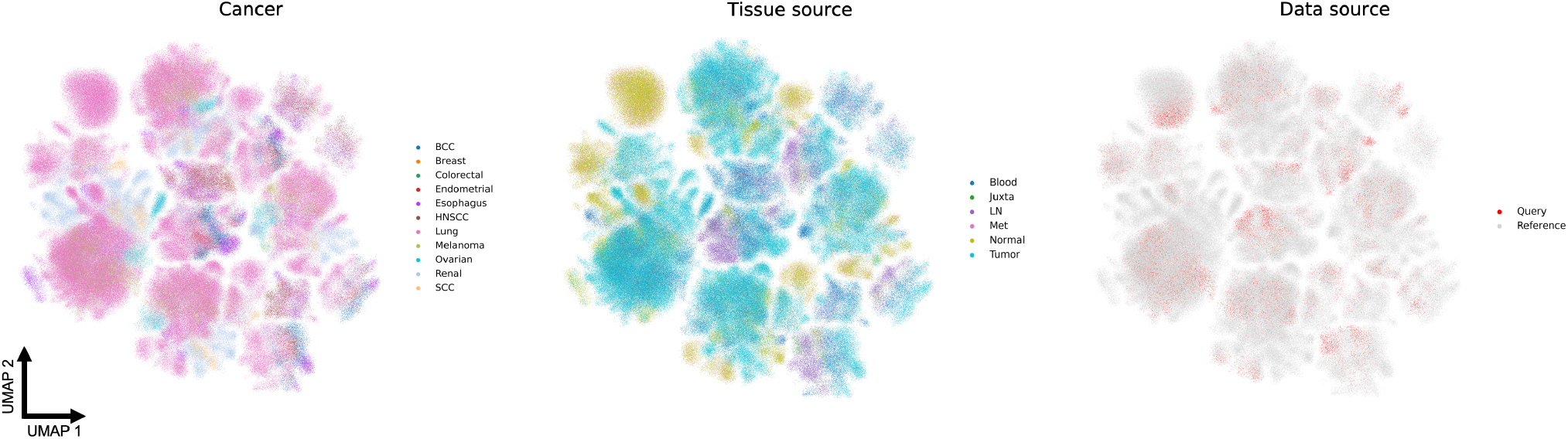
Embedding of the TIL dataset without scArches. UMAP visualizations of the joint embedding colored by cancer tissue, tissue type, and query vs. reference set.

**Supplementary Data 1** | **Benchmarking results**. Quantitative evaluation and statistical analysis of the models’ performance on the 10x dataset (both donors) and the Fischer dataset between the unimodal and multimodal models, and *tessa*.

**Supplementary Data 2** | **Cluster selection on the SARS-CoV-2 dataset**. Statistical analysis of the IFN response score across clusters and similarity of transcriptome and TCRs of CD8^+^ T effector cells within selected clusters.

**Supplementary Data 3** | **HLA annotation for the SARS-CoV-2 dataset**. Information about HLA type for each donor in the SARS-CoV-2 dataset.

**Supplementary Data 4** | **IEDB-based Epitope Query for selected clusters of the SARS-CoV-2 dataset**. Pairs of TCRs and epitopes from selected clusters of the SARS-CoV-2 dataset derived from a database query to the *IEDB*.

**Supplementary Data 5** | **DEG analysis of CD8**^**+**^ **T effector cells in the SARS-CoV-2 dataset**. Log-fold change, p-values, and adjusted p-values of the genes comparing disease-specific and bystander clusters in the SARS-CoV-2 dataset.

## Notes

### Summary of Updates

Revision includes the following changes: - extended benchmark - application showcase on SARS-CoV-2 dataset - run time analysis and integration capability over multiple studies

